# Common *cis*-eQTLs do not explain chromosome 21 gene expression deviations from the ploidy expectation in Trisomy 21

**DOI:** 10.64898/2026.10.10.758221

**Authors:** Lucas A. Gillenwater, Mary A. Allen, Milton Pividori, Casey S. Greene, Marc Subirana-Granés

## Abstract

Down syndrome (DS) results from trisomy 21 (T21), the triplication of human chromosome 21 (HSA21). On average, genes on HSA21 are overexpressed at the expected 1.5-fold increase in people with DS. A small subset of HSA21 genes is expressed closer to euploid levels. Could local genetic regulatory variation explain deviations from the expected dosage? Using the largest DS-specific transcriptomic dataset to date from the Human Trisome Project (HTP) and common GTEx expression quantitative trait loci (eQTLs), we tested whether local genetic regulatory variation contributes to gene-specific buffering or amplification of HSA21 dosage effects in DS.

We compared the bulk RNA-seq profiles of protein-coding genes, lncRNAs, and pseudogenes from whole-blood samples of 274 participants with T21 and 80 euploid (D21) participants. We filtered out genes with low read counts or high genomic repeat content and applied ploidy correction to the differential expression analysis. We found 13 genes that differed significantly from the expected fold change. We cross-referenced GTEx whole-blood eQTLs (*cis* window ±1 Mb) with the 13 deviating genes. A common *cis*-eQTL was detected for 8 of the 10 testable genes. However, 59 of 99 HSA21 genes that had the expected 1.5-fold change had significant *cis*-eQTLs. Additionally, T21 participants carry the eQTL alleles at similar frequencies to GTEx donors. Therefore, *cis*-eQTLs explain differences between people with T21, not the difference between T21 and D21.

Overall, common *cis*-regulatory variation acts on HSA21 genes in T21 as it does in euploid blood, but it does not explain which genes deviate from the ploidy expectation.

## Introduction

The triplication of human chromosome 21 (HSA21), trisomy 21 (T21), causes Down syndrome (DS). Multiple studies report that genes on HSA21 are expressed, on average, at the expected 1.5-fold change when compared to people with euploid karyotypes (disomy 21, D21) [1,2,3,4]. A small subset of HSA21 genes is expressed closer to euploid levels. Potential causes for lower-than-expected HSA21 gene expression include confounding cell-type proportions, genetic effects, and compensatory regulation. This study assesses the genetic effect hypothesis while controlling for cell-type proportions.

Previous researchers have noted that differences in genetic composition affect clinical and molecular phenotypes in DS. Rachubinski et al. hypothesized that genetic variations beyond T21 may explain the development of autism spectrum disorder in some individuals with DS [5]. Valentini et al. assessed genomic effects with a genome-wide association study (GWAS) in a cohort of 84 children with DS and found that the *PNPLA3* variant rs738409 on HSA22 was associated with liver steatosis [6]. A review by Antonarakis et al. noted that ∼80% of T21 conceptuses are lost during pregnancy and inferred that a specific combination of genetic variants allows T21 conceptuses to survive to term [7]. In 388 live-born individuals with DS, Popadin et al. found signatures of embryonic selection: a deficit of deleterious variants on HSA21 and reduced transcriptome-wide variation in the expression of highly constrained genes [8].

Hunter et al. tested whether the genotype of common expression quantitative trait loci (eQTLs) explained lower-than-expected gene expression of HSA21 genes in a child with DS compared to family members (i.e., brother, mother, and father, all euploid for HSA21) [9]. The authors introduced a methodology to control for common factors that bias differential expression studies in DS research. Specifically, the methodology accounted for genes with no reads, low gene expression, high genomic repeats, and karyotype. Overall, the analysis confirmed that the majority of HSA21 genes show the expected 1.5-fold change. There were 5 genes that fell below the expected fold change. Of those 5 genes, 3 had common eQTLs in the general population. Genotyping confirmed that the allele dosage at those eQTLs explained the lower-than-expected gene expression in the individual with DS.

Here we extend the methodology of Hunter et al. to the population level. We assess how *cis*-eQTLs on HSA21 affect whole-blood gene expression in a large cohort of individuals with and without DS. We account for triploidy and cell-type composition in the differential expression analysis. We perform ploidy-aware genotyping of HSA21 in people with T21 to accurately represent allele counts. We compare the sample genotypes against common eQTLs found in whole blood of disomic individuals in the GTEx cohort. Overall, common *cis*-eQTLs affect HSA21 gene expression in people with T21 as they do in D21 individuals, but they do not explain which genes deviate from the expected 1.5-fold change. The genotype of one person can account for that person’s deviations, as in Hunter et al., but common *cis*-regulatory variation does not explain the deviation at the population level.

## Methods

### Study consent

As part of the INCLUDE project, the Linda Crnic Institute for Down Syndrome at the University of Colorado Anschutz Medical Campus enrolled participants under a study protocol approved by the Colorado Multiple Institutional Review Board (COMIRB #15-2170). Human Trisome Project (HTP) researchers at the Linda Crnic Institute for Down Syndrome obtained written informed consent from all participants over 7 years old, as allowed by their cognitive ability, or from their legally authorized representative. Regardless of study participant age or cognitive ability, researchers explained study procedures to the participant. HTP researchers recorded age, sex, and body mass index (BMI) during the visit.

### Whole-blood transcriptome quantification

Waugh et al. [1] describe the process of blood collection, processing, and quantification in detail. Briefly, researchers collected peripheral blood in PAXgene RNA tubes (QIAGEN), performed poly-A+ RNA enrichment, and used the Illumina NovaSeq 6000 (Novogene) for paired-end sequencing before additional bioinformatic processing. The libraries were globin-depleted and strand-specific (NEBNext Ultra II Directional), and gene-level counts were obtained by strand with HTSeq-count against the GENCODE v33 annotation (GEO GSE190125).

### Mass cytometry of white blood cells

For a detailed account of the quantification of white blood cells, see Galbraith et al. [2]. Whole-blood samples underwent red blood cell lysis and white blood cell fixation, followed by barcoding with a Cell-ID 20-Plex Pd kit and pooling. Pooled samples were stained with antibodies, with surface marker staining performed before permeabilization and intracellular transcription factor/phospho-epitope staining performed after. Cells were then labeled with Cell-ID Intercalator-Ir and quantified on a Helios mass cytometer. Batched FCS files were demultiplexed and individual samples were normalized across batches.

### Whole genome sequencing

Whole genome sequencing of Human Trisome Project participants was performed at the Broad Institute as part of the NIH INCLUDE and Gabriella Miller Kids First projects, and the cohort is described by Feldman et al. [10]. PCR-free libraries were prepared from genomic DNA sheared to approximately 385 bp using the KAPA Hyper Prep kit without amplification, and sequenced on an Illumina NovaSeq 6000 (S4 flow cells) to a target read depth of ∼30x coverage to produce 151 bp paired-end reads.

Reads were aligned to the GRCh38 reference genome (Homo_sapiens_assembly38) with BWA-MEM v0.7.15, duplicate-marked with Picard v2.20.4, and base quality score recalibrated with GATK v4.0.10.1 using the Broad WholeGenomeGermlineSingleSample workflow. These analysis-ready CRAM files were used as the input for variant calling, without further preprocessing.

### Ploidy-aware variant calling on HSA21

Standard germline genotyping workflows assume a diploid genome and therefore misassign allele dosage on HSA21 in individuals with T21; for this reason, chromosome 21 was excluded from the association analyses of Feldman et al. [10]. We therefore re-called HSA21 variants directly from the aligned CRAM files with a ploidy-aware caller. Variants on HSA21 were called in 433 Human Trisome Project participants with whole genome sequencing data, comprising 419 participants with T21 and 14 D21 participants. Variants were called against the same GRCh38 reference with Octopus v0.7.4 [11], using the individual caller on each sample independently and restricting calling to HSA21 (–regions chr21). The ploidy of HSA21 was set to 3 for participants with T21 (–contig-ploidies chr21=3), whereas D21 samples were called at the default ploidy of 2. All other Octopus parameters were left at their default values.

Calls were filtered with the default Octopus hard-filter expression (QUAL < 10, MQ < 10, MP < 10, AD < 1, AF < 0.01, AFB > 0.25, SB > 0.98, BQ < 15, DP < 1, ADP < 1), and only records with FILTER = PASS were retained using BCFtools v1.21 [12]. Filtering retained approximately 85% of raw calls, corresponding to a median of 81,667 variants per T21 sample (range, 59,473 to 104,854) and 66,238 variants per D21 sample (range, 63,054 to 80,735).

Within each group, PASS-filtered per-sample calls were merged with bcftools merge, keeping distinct alternate alleles at the same position as separate records (–merge none). Samples with no call at a merged site were assigned the homozygous reference genotype (–missing-to-ref; 0/0/0 in T21, 0/0 in D21). The samples were not jointly genotyped, so a missing call could not be distinguished from a homozygous reference genotype. The missingness may have lowered alternate allele frequencies at sites with incomplete calls. We therefore checked the T21 allele frequencies against those of GTEx whole-blood donors for a systematic offset (see *cis*-eQTL comparisons). The resulting call sets comprised 688,787 variant records in the T21 group and 189,474 in the D21 group. Of these, 529,386 T21 and 142,792 D21 records contained a single nucleotide variant (SNV), and 160,468 T21 and 46,845 D21 records contained an indel; a multiallelic record can contain both, so the two counts overlap. The workflow was implemented in Snakemake v9.16.3 [13] and run on the Alpine high-performance computing cluster (University of Colorado Boulder Research Computing).

### Cohort filtering

To assess genetic effects, we required samples to have matched whole-blood bulk RNA sequencing counts, whole genome sequencing, cell-type composition, and all demographic and technical data (i.e., age, sex, BMI, and sample source) (Figure S1A). Additionally, while ∼95% of individuals with DS have full T21, translocation accounts for ∼5% and mosaicism for ∼2% of cases [7]. Mosaicism affects expression levels (Figure S1B). Therefore, we removed samples with mosaic T21 (9 samples). We kept samples with translocation DS, whose HSA21 expression was similar to that of samples with full T21 (Figure S1B). The 274 T21 participants in the analysis cohort comprised 235 with full T21, 7 with translocation DS, and 32 whose karyotype subtype was not recorded (we therefore assume full T21).

### Ploidy-aware differential expression analysis

Generally, we followed the methods of Hunter et al. [9]. We compared DESeq2 [14] fits using default parameters against fits that set the expected fold change to 1.5 for HSA21 genes in T21 samples and to 1 for all other gene-karyotype combinations. We excluded HSA21 from the size-factor (library normalization) calculation so that HSA21’s own inflated counts do not bias normalization for the rest of the genome. As a negative control, we assessed the change in distribution for HSA22 genes between the run with default parameters and the ploidy-adjusted run.

The model included age, sex, BMI, and sample source as covariates since those variables were associated with karyotype (Table 1). In addition, cell-type proportions affect gene expression (Figure S1C). Therefore, we included 19 cell-type proportions as covariates in the model. P-values were adjusted for multiple testing across genes using the Benjamini-Hochberg procedure [15].

**Table 1.**
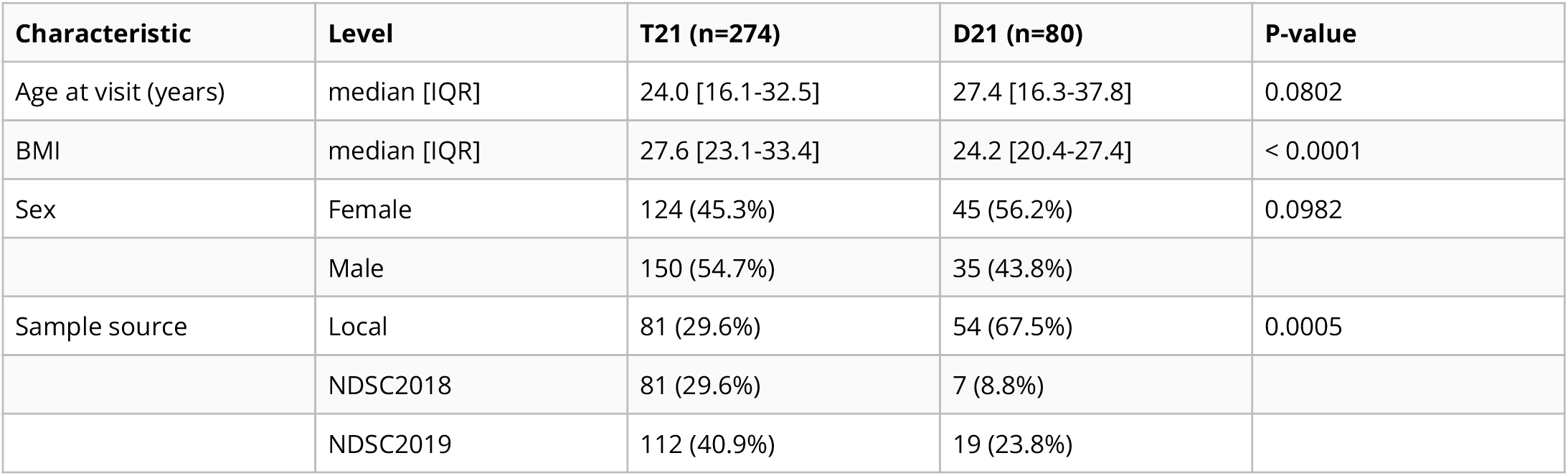
Analysis cohort assembly and baseline characteristics, stratified by karyotype. P-values compare T21 and D21 participants: Wilcoxon rank-sum test for age and BMI, and Fisher’s exact test for sex and sample source (for sample source, which has three levels, the p-value was simulated with 2,000 replicates).

We restricted the analysis of HSA21 genes to protein-coding, lncRNA, and pseudogenes since those are the gene biotypes covered by the GTEx eQTL data. We excluded genes with low average read counts (baseMean < 30) or that fall in genomic regions with high repeat content that cannot be reliably quantified. A gene was classified as deviating from the 1.5-fold ploidy expectation if its ploidy-corrected fold change differed from 1.5 by at least 33% (adjusted p < 0.01). Deviating genes were further stratified into tier 1 (≥50% deviation from the 1.5-fold expectation) and tier 2 (33–50% deviation). The filters and classification of each targeted HSA21 gene are given in Supplementary Table S1.

### *cis*-eQTL association testing

For each deviating gene, we retrieved all variants within ±1 Mb that GTEx v10 annotates as at least nominally associated (GTEx p ≤ 10^−4^) with that gene’s expression in whole blood. The minor allele refers to the GTEx whole-blood minor allele frequency and checked against gnomAD v4.1 (Supplementary Table S2). We regressed expression against genotype dosage (0-3 copies of the alternate minor allele) within T21 participants, with the covariates of the primary analysis (age, sex, BMI, sample source, and 19 cell-type proportions) regressed out of both expression and genotype dosage and retained each gene’s strongest-associated variant. For each gene, we report the effect per copy of whichever allele, minor or major, correlates with the direction of the fold-change between T21 and D21 participants. Statistical significance was assessed by a gene-level permutation test where the covariate-adjusted expression values were shuffled across participants 1,000 times, the same variant search was repeated on each shuffle, and the fraction of shuffles producing an association at least as strong as the observed one gave a gene-level permutation p-value. Permutation p-values were adjusted for multiple testing across genes using the Benjamini-Hochberg procedure [15]. A gene’s *cis*-eQTL was considered detected at q < 0.05.

### *cis*-eQTL comparisons

Under an additive model, each copy of HSA21 contributes to gene expression. The T21/D21 ratio is then 1.5 × (1 + *r* × *p*_T21_) / (1 + *r* × *p*_D21_), where *r* is the output of the alternate allele relative to the reference allele, and *p* is the frequency of the alternate allele in each group. If the allele has the same frequency in both populations, the ratio is 1.5.

The cohort had 14 D21 participants with WGS, only 2 of which had matching transcriptomic data. Therefore, we estimated D21 allele frequencies from the GTEx whole-blood donors. The T21 participants contribute 822 alleles (3 each), and the 800 GTEx whole-blood donors contribute 1,600 (Supplementary Table S2 also gives the frequencies in gnomAD v4.1). Confidence intervals for the difference assume binomial sampling in both groups. Therefore, a T21 participant’s three copies are assumed independent, although two come from the same parent. We did not have parental WGS and therefore could not determine which parent experienced nondisjunction prior to mating.

For each tested gene’s best variant, we obtained *r* from the within-T21 slope (*r* = 2^3*b*^ − 1, where *b* is the slope in log_2_ units per copy) and took the two allele frequencies from the T21 participants and from GTEx. The ratio between T21 and D21 reveals the shift from 1.5 that the variant produces at the observed frequencies. We then artificially set the T21 frequency to 0 and to 1 to find the largest shift the variant could produce at any frequency. We solved the ratio for the T21 frequency that would produce the entire deviation. We also report how much of the expression variance among T21 participants the best variant explains.

We compared the deviating genes with the HSA21 genes at the expected dosage. We applied the identical test (same variant selection, participants, covariates, and 1,000 permutations) to every expected-dosage gene with a GTEx variant to test, adjusting for multiple testing across those genes. Detection depends on the strength of an eQTL, so we compared the deviating genes with expected-dosage genes whose eQTLs are of similar strength in GTEx. Among the expected-dosage genes that are GTEx whole-blood eGenes (i.e., have an eQTL at q < 0.05), we fit a logistic regression of detection on the log2 of the absolute GTEx allelic fold change (aFC) [16] at the GTEx lead variant, and compared the observed number of detected deviating genes with the distribution of the sum of their predicted probabilities. To compare effect sizes, we estimated the aFC in T21 by least squares, modeling each of the three HSA21 copies as contributing its own expression and fitting the covariates jointly. Each dataset’s effect was taken at its own strongest variant, so that both carry the same selection: the T21 aFC at the variant selected by the within-T21 test, signed by the GTEx direction at that variant, was divided by the absolute GTEx aFC at the GTEx lead variant. We compared this ratio between the deviating genes and the expected-dosage eGenes whose absolute GTEx aFC lay within the deviating genes’ range, using a Wilcoxon rank-sum test. To estimate the power of the test for each tested deviating gene, we permuted its expression across participants, added an allelic effect at its GTEx lead variant under the three-copy model, and repeated the gene-level test, 200 times for each of a grid of effect sizes. A simulation counted as a detection at gene-level permutation p < 0.05, and power at the gene’s GTEx aFC was interpolated from the grid. As a negative control, we regressed the expression of every assessable HSA21 gene on every common HSA21 variant at least 5 Mb from its transcription start site, where no *cis* effect is expected, and measured the genomic inflation factor (Supplement B, Figure S2).

### Expression of the HSA21 genes in blood

To check that the deviating genes are expressed in blood, we took the median expression of each HSA21 gene in 68 tissues from GTEx v10, in transcripts per million (TPM) [17]. We compared the whole-blood TPM of the deviating and expected-dosage genes with a Wilcoxon rank-sum test. We also placed each deviating gene at its percentile among the expected-dosage genes.

To relate expression to eQTL detection, we compared whole-blood TPM between GTEx eGenes and non-eGenes, and between genes with and without a *cis*-eQTL detected in T21, with Wilcoxon rank-sum tests. We ranked each deviating gene’s whole-blood TPM among its 68 GTEx tissues to assess the typical representation in blood.

We also compared expression in the T21 participants with GTEx whole blood. We converted the participants’ read counts to TPM, using the union of exons in the GENCODE v44 basic annotation as gene length [18]. Since the HTP libraries are globin-depleted and the GTEx libraries are not, we measured the overall offset between the two datasets on the non-HSA21 genes.

### Sensitivity analysis

To quantify the effect of covariate adjustment and mosaic exclusion, we compared the adjusted results to the outputs from the identical pipeline without covariate adjustment and without excluding mosaic participants (Figures S1D and S3). Additionally, we refit the adjusted model with an interaction between karyotype and each of the 19 cell fractions. The fractions were centered, so the karyotype effect is the T21 effect at the cohort-mean composition. The interaction terms widened the standard error of the karyotype effect by a median of 1.76-fold for HSA21 genes, so we compared effect estimates rather than significance across the models (Supplement C, Table S3).

## Results

### Most chromosome 21 genes show a gene-dosage-dependent increase in expression

To determine whether HSA21 genes in whole blood from people with T21 are expressed in proportion to their extra copy, we compared transcript levels between T21 and D21 participants before and after ploidy correction.

The HTP RNA-seq cohort consisted of 399 participants, 304 with T21 and 95 D21. Filtering removed samples missing cell-type composition (22) or BMI metadata (12), and samples with mosaic karyotypes for HSA21 (9). We also removed samples from T21 participants who did not have WGS data (2). We compared whole-blood bulk RNA transcript counts between 274 T21 samples and 80 D21 samples, a total of 354 samples. T21 participants had significantly higher body mass index (BMI) than D21 participants (Table 1). The D21 participants were older than the T21 participants (median 27.4 vs. 24.0 years), but the difference was not significant. The sex composition did not significantly differ by karyotype, though a higher proportion of T21 participants were male. Though processed at the same facility, the samples were collected at 3 different locations.

We assessed the effects of ploidy correction on the differential expression analysis. Of the 318 HSA21 genes, 153 had baseMean coverage 30 and were not in high-repeat regions. Before ploidy correction, statistically significant (adjusted p < 0.01) fold changes of at least 1.33 in either direction were observed in 76% (117) of the HSA21 genes passing thresholds. After ploidy correction, 13 genes (8.5%) still deviated from the expected 1.5-fold change by at least 33% in either direction, at adjusted p < 0.01 (Table S4 and Figures 1A–B and S4). The deviations of *OLIG2, BACE2, OLIG1*, and *RUNX1* were the most robust to the choice of covariate model (Supplement C). Per-gene statistics for all 318 genes are given in Supplementary Table S1. The magnitude of the correction matches the ploidy expectation. Across the 318 HSA21 genes, the median log2 fold change shifted from 0.593 (close to log2(1.5) = 0.585) before correction to 0.008 after correction. Chromosome 22 showed no shift, confirming that the correction is specific to HSA21 (Figure 1C–D).

**Figure 1:**
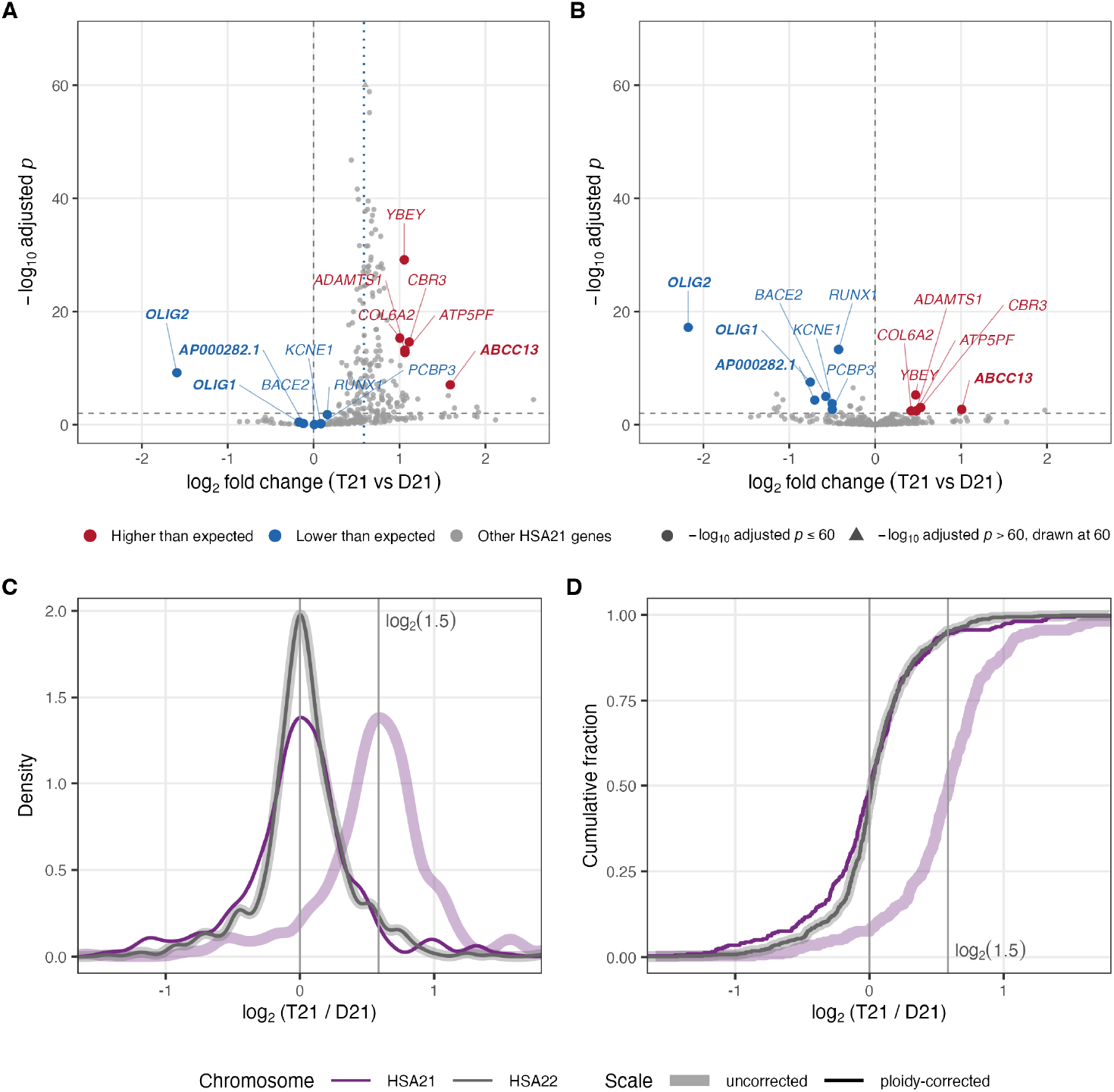
Differential expression of HSA21 genes between T21 and D21 participants, before and after ploidy correction. (A, B) The 318 targeted HSA21 genes before (A) and after (B) ploidy correction. Each point is one gene, plotted by its log _2_ fold change and the −log_10_ of its adjusted p-value. The dotted line in A marks the 1.5-fold expectation (log _2_ 1.5 = 0.585), and the dashed horizontal line marks adjusted p = 0.01. The 13 deviating genes are labeled and colored by direction (red, higher than expected; blue, lower than expected), with tier 1 genes in bold; other HSA21 genes are grey. (C) Density and (D) empirical cumulative distribution of the log_2_ fold change (T21 vs D21) for the 318 targeted HSA21 genes (purple) and 650 HSA22 genes (grey), on the uncorrected (light) and ploidy-corrected (dark) scales. Vertical lines mark 0 and log _2_ 1.5.

### Common *cis*-eQTLs affect gene expression in T21, but do not explain deviations in gene expression

We corrected for ploidy in the differential expression analysis to find HSA21 genes that deviated from the expected dosage. A departure could reflect regulation that offsets or amplifies the extra copy, or common *cis*-regulatory alleles that differ in frequency between people with and without T21. To test the genetic explanation, we compared the ploidy-aware genotypes and covariate-adjusted whole-blood gene expression of 274 T21 participants to common whole-blood eQTLs.

Three of the 13 deviating genes had no variant to test: GTEx tested *ATP5PF* and *RUNX1* in whole blood but found no eQTL (q = 0.22 and 0.21) [17], and GTEx did not test *AP000282*.*1* in whole blood (the GTEx gene model excluded the exon it shares with *OLIG1*; Supplement E). Of the remaining 10 testable genes, 8 have a detected *cis*-eQTL (gene-level permutation p < 0.001 for six, p = 0.002 for *KCNE1*, and p = 0.027 for *ABCC13*) and 2 do not (*BACE2*, p = 0.30; *OLIG2*, p = 0.40) (Table 2, Figure 2A–C). The expression of *ABCC13* (q = 0.034) is also associated with HSA21 genotype far from its own locus (Supplement B). All 13 deviating genes sit on the q arm, between 14.2 and 46.3 Mb (Figure 2D). The deviating genes are measurably expressed in blood (Supplement E, Figure S5A). Expression level in GTEx did not correspond with eQTL detection (Figure S5B). *OLIG2* and *BACE2* are among the least expressed in GTEx whole blood (0.27 and 0.71 TPM), yet GTEx detects strong whole-blood eQTLs for both (q = 1.2 × 10^−13^ and 6.2 × 10^−16^). Allele frequencies for every tested variant, in GTEx whole blood, in gnomAD, and in this cohort, are given in Supplementary Table S2.

**Table 2.**
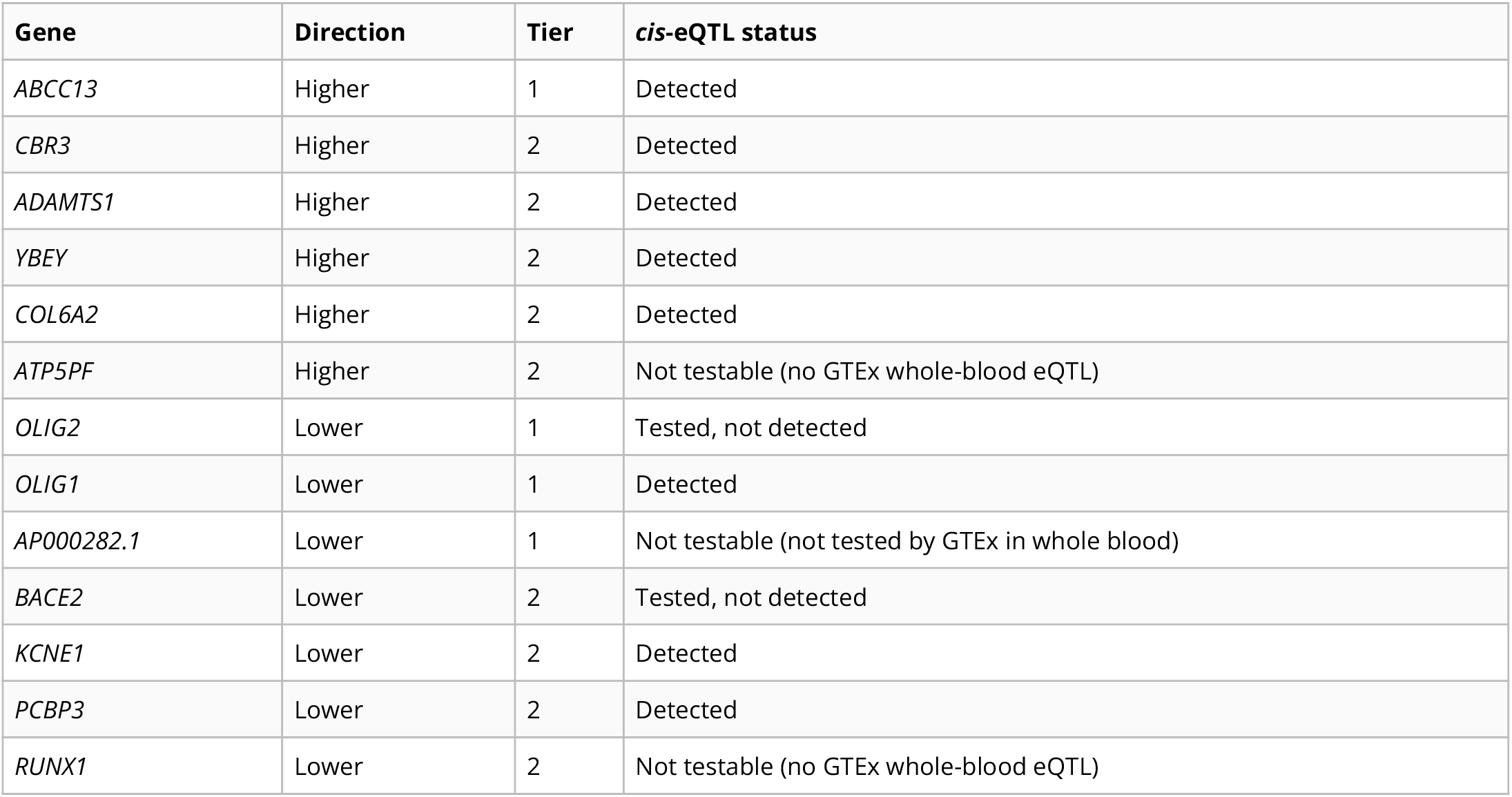
*cis*-eQTL status of the 13 genes that deviate from the ploidy expectation in the primary (adjusted) analysis.

| Gene | Direction | Tier | *cis*-eQTL status |
| --- | --- | --- | --- |
| *ABCC13* | Higher | 1 | Detected |
| *CBR3* | Higher | 2 | Detected |
| *ADAMTS1* | Higher | 2 | Detected |
| *YBEY* | Higher | 2 | Detected |
| *COL6A2* | Higher | 2 | Detected |
| *ATP5PF* | Higher | 2 | Not testable (no GTEx whole-blood eQTL) |
| *OLIG2* | Lower | 1 | Tested, not detected |
| *OLIG1* | Lower | 1 | Detected |
| *AP000282.1* | Lower | 1 | Not testable (not tested by GTEx in whole blood) |
| *BACE2* | Lower | 2 | Tested, not detected |
| *KCNE1* | Lower | 2 | Detected |
| *PCBP3* | Lower | 2 | Detected |
| *RUNX1* | Lower | 2 | Not testable (no GTEx whole-blood eQTL) |

**Figure 2:**
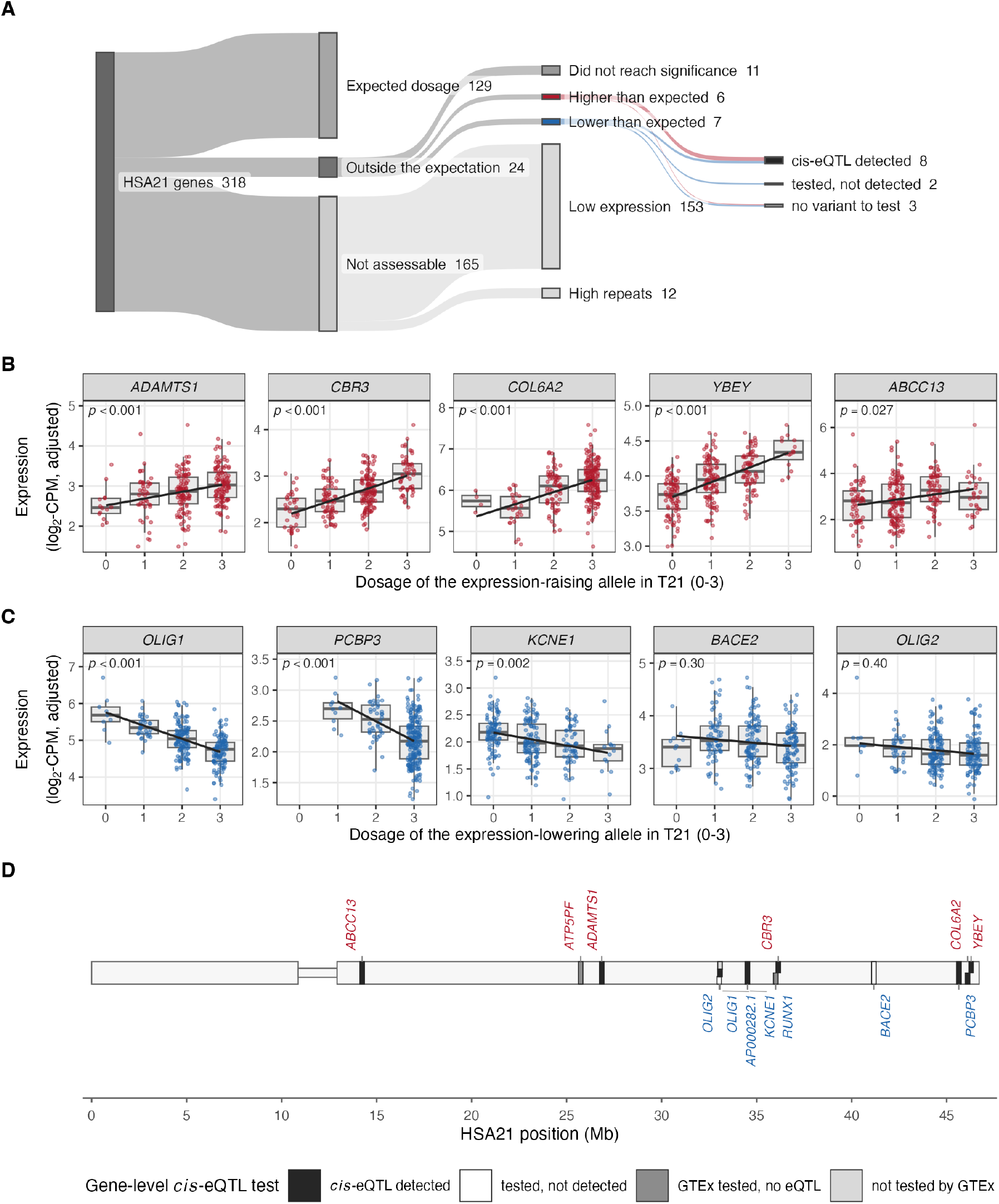
Classification of the 318 targeted HSA21 genes, expression by genotype, and positions of the deviating genes. (A) Sankey plot of every gene from the full set through the dosage classification. (B, C) Expression against genotype at the best variant for the 10 testable deviating genes, in the 274 T21 participants. Each panel plots covariate adjusted expression against the number of copies (0 to 3) of the allele that GTEx links to the gene’s direction of deviation. Boxes show the median and interquartile range, and the line is the within-T21 fit. Each panel gives the gene-level permutation p-value of the *cis*-eQTL test (1,000 permutations, with the variant search repeated on each). (D) Chromosome 21 (GRCh38) with a band at the transcription start site of each deviating gene, colored by *cis*-eQTL outcome: detected (black), tested without detection (white), or no variant to test, either because GTEx tested the gene in whole blood and found no eQTL (dark grey; *ATP5PF, RUNX1*) or because GTEx did not test it (light grey; *AP000282*.*1*).

### The detected *cis*-eQTLs cannot account for the deviations in gene expression

To ask whether a detected *cis*-eQTL is more common among the deviating genes than among HSA21 genes at the expected dosage, we applied the identical test to the 99 testable expected-dosage genes, of which 59 (60%) had a detected *cis*-eQTL. The effects in T21 were similar in size relative to the GTEx effect. The median T21 allelic fold change was 0.73 for the deviating genes and 0.76 for the 49 expected-dosage eGenes with GTEx effects in the same range (Wilcoxon p = 0.64; Figure 3A). As a negative control, the expression of the assessable HSA21 genes regressed on common HSA21 variants at least 5 Mb away showed no inflation beyond its permutation range (λ = 1.020; Supplement B). *BACE2* and *OLIG2* were not missed for lack of power. Effect spike-in tests detected effects at power > .8 for all but 2 of the deviating genes, *ADAMTS1* and *ABCC13* 3B).

**Figure 3:**
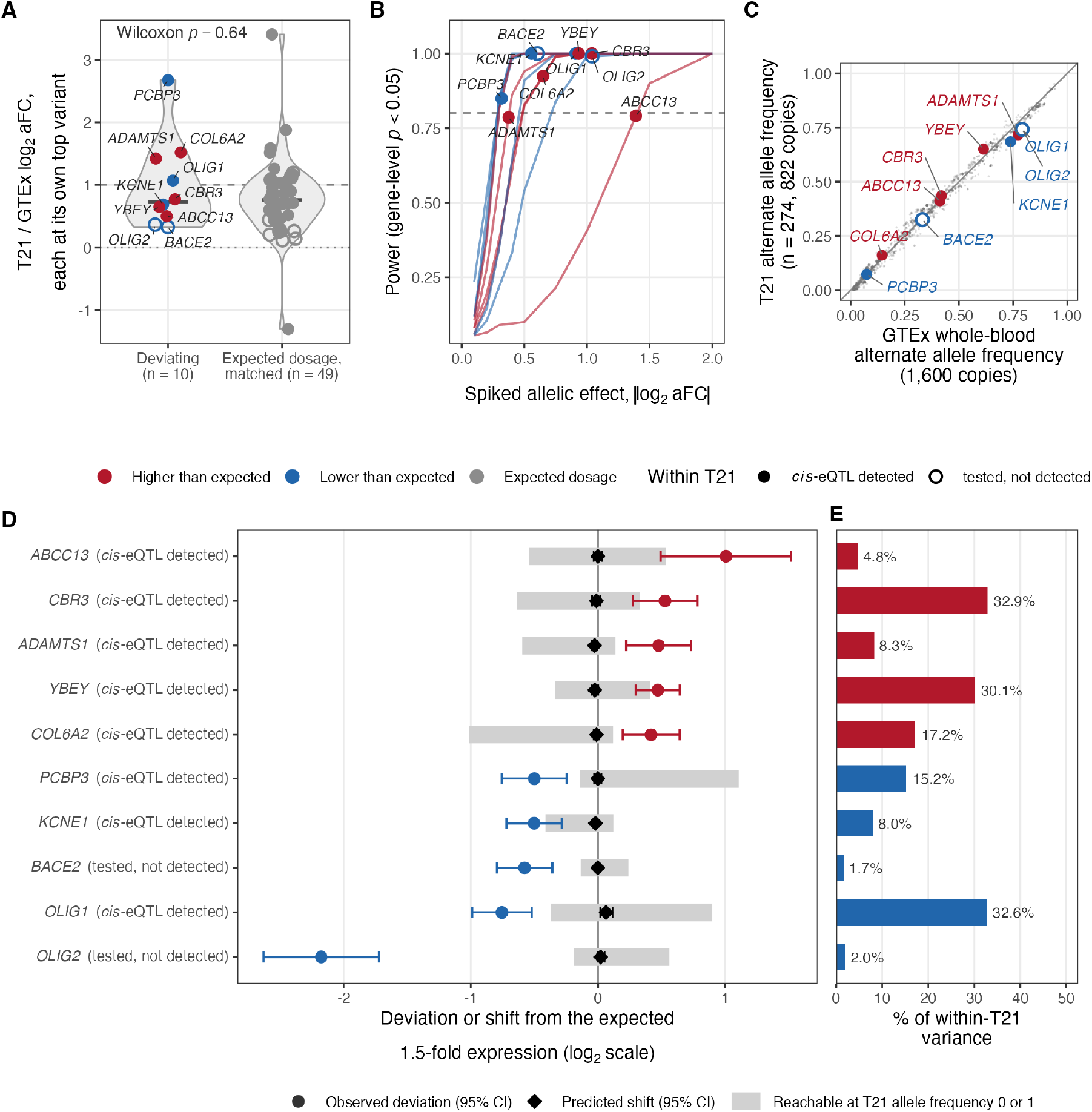
Effect size and power of the *cis*-eQTL test, and how much of each deviation a *cis*-eQTL could produce. (A) For each gene, the T21 allelic fold change at its strongest variant in T21 relative to the GTEx whole-blood allelic fold change at the GTEx lead variant, for the 10 testable deviating genes and the 49 expected-dosage GTEx eGenes whose GTEx effect lies in the same range (|log_2_ aFC| 0.32 to 1.39). The dashed line marks an effect equal to GTEx’s and the dotted line none. Deviating genes are colored by direction and expected-dosage genes are grey. (B) Power of the gene-level test for each tested deviating gene, from 200 simulations per effect size with the gene’s own genotypes, variant set, and expression noise. Points mark each gene’s GTEx effect, and the dashed line marks 80%. (C) Alternate allele frequency of the 2,076 tested *cis* variants in the 274 T21 participants (822 copies) against the 800 GTEx whole-blood donors (1,600 copies), with the deviating genes’ best variants labeled (filled, *cis*-eQTL detected in T21. (D) For each tested deviating gene, the observed deviation of log_2_(T21/D21) from log_2_ 1.5 (point, 95% CI, colored by direction as in A), the shift predicted by the observed frequency difference at its best variant (diamond, 95% CI), and the largest shift possible, with the alternate allele on every T21 copy or on none, taken at the 95% confidence edge of the slope (grey bar). (E) Fraction of expression variance among T21 participants explained by the best variant.

The T21 participants carried the tested alleles at frequencies that did not differ systematically from those of the 800 GTEx whole-blood donors. Across the 2,076 tested *cis* variants, the median difference was −0.0005 (Figure 3C). At each tested gene’s best variant, the observed frequency difference predicted between −8% and +4% of the deviation (Figure 3D). Even if T21 participants carried the allele on every copy or on none, the best variant could not produce the observed deviation. The best variants explain 2% to 33% of the expression variance among T21 participants (Figure 3E).

In summary, after ploidy correction most HSA21 genes showed the expected increase with gene dosage, and 13 deviated. A common *cis*-eQTL was detected for 8 of the 10 testable deviating genes, about as often as for HSA21 genes at the expected dosage with eQTLs of the same GTEx strength, and with similar effect sizes. The T21 participants have allele frequencies that do not differ systematically from those of GTEx donors. *cis*-eQTLs explain differences between people with T21, not the difference between T21 and D21. Our work does not support the genetic hypothesis for expression deviations for any of the 13 genes outside of the 1.5-fold difference, including the 2 tested without a detected eQTL (*BACE2, OLIG2*) and the 3 that could not be tested (*ATP5PF, RUNX1, AP000282*.*1*) (Figure 2A).

## Discussion

Trisomy 21 increases the expression of most HSA21 genes by the expected 1.5-fold, yet there is a subset of HSA21 genes with expression more similar to that of disomic individuals [4,9]. Lower-or higher-than-expected gene expression may reflect genetic variation across the population, compensatory regulation, or technical artifacts. Hunter et al. tested the genetic regulation hypothesis within a family cohort consisting of one T21 child, one D21 child, and the two D21 parents [9]. No one had yet assessed the genetic variation hypothesis for deviating gene expression using transcriptomic and WGS profiles from a large, population-level cohort of people with DS. Here we demonstrate that common *cis*-eQTLs affect HSA21 gene expression in people with T21. However, the common *cis*-regulatory variation does not explain gene expression deviation from the expected 1.5-fold difference at the population level.

Hunter et al. found that only 5 genes had expression levels similar to D21 levels after considering coverage, high-repeat regions, and proper ploidy in the differential expression analysis. Through cross-referencing common *cis*-eQTLs from GTEx whole-blood profiles, the authors found that the genotype of the individual with DS explained the variance in *CLIC6, ITSN1*, and *C2CD2*, but not for *LINC02246* or *BRWD1*. In the present study, we did not find the same 5 deviating genes that Hunter et al. found. *ITSN1, C2CD2*, and *BRWD1* are expressed in whole blood and sit at the expected dosage. *CLIC6* and *LINC02246* had low coverage.

In a cohort of 4, where one individual had T21, every expression difference between that person and the others reflects that person’s genotype, including their eQTL alleles [9]. A different set of four people would have different alleles and would show eQTL-driven differences at a different set of HSA21 genes. A population-level study asks whether eQTL effects produce a systematic difference between people with and without T21. The present study evaluated 354 whole-blood profiles and found that the tested alleles occur at similar frequencies in people with T21 and in euploid GTEx donors (Figure 3C). Both studies show genetic effects on gene expression in people with DS (Figure 3A), but at the population level eQTLs did not explain the deviations from the expected 1.5-fold change (Figure 3D). Expression differences in a small sample reflect the alleles of the people sampled rather than persistent differences in people with T21.

We demonstrated that the majority of HSA21 genes show the expected 1.5-fold change in gene expression if we filter out genes in high-repeat regions and genes with low coverage, and adjust the differential expression pipeline for sample ploidy. We tested for deviation both above and below the expected dosage. The compensation hypothesis predicts that genes are downregulated toward euploid levels, whereas eQTLs can raise or lower expression. Of the 10 deviating genes with a GTEx variant to test, 5 were higher than expected and 5 lower, and 8 had a detected *cis*-eQTL.

The two genes that did not have *cis*-eQTLs were *BACE2* and *OLIG2. BACE2* codes for a homolog of β-secretase BACE1, a protein involved in amyloid precursor processing and associated with Alzheimer’s disease in people with DS [19]. *OLIG2* codes for a transcription factor involved in mammalian central nervous system development [20]. Both were found to have lower expression in the population than expected, closer to the euploid level. Both are known for their roles in brain tissue and are expressed at low levels in blood, but GTEx detects strong whole-blood eQTLs for both, and the test had full power at their GTEx effect sizes (Figure 3B; Supplement E). Their T21 effects relative to GTEx (0.33 and 0.36) are within the range of matched expected-dosage eGenes in GTEx that were also not detected, and their deviations are among the most robust to the choice of covariate model (Supplement C).

Overall, the genomic hypothesis does not explain deviating genes in this cohort. A *cis*-eQTL was detected for 60% of the expected-dosage HSA21 genes that could be tested, so the deviating genes were not unique. The eQTL effects in T21 had the same direction and a similar size as in GTEx (Figure 3A). The mechanism for the deviation of these genes therefore remains open.

There are multiple other factors that may explain deviations in gene expression. Candidate non-genetic mechanisms for lower-than-expected transcript counts include autoregulatory feedback loops, post-transcriptional silencing by triplicated HSA21 miRNAs, alternative splicing into unstable isoforms, and chromatin remodeling. In either direction, an HSA21 gene may also deviate because it lies downstream of pathways that other triplicated HSA21 genes misregulate. Three of the 7 genes lower than expected encode transcription factors (*OLIG1, OLIG2*, and *RUNX1*), a class in which autoregulation is common [21]. Negative autoregulation would pull a gene’s expression back toward the euploid level. For example, Laufer et al. proposed that hypermethylation of an enhancer within *RUNX1* by DNMT3A and DNMT3L attenuates RUNX1 overexpression by the time of birth in people with DS [22]. However, in this study *OLIG2* was expressed at one-third of the D21 level in people with DS, and therefore needs an additional explanation.

Six of the 13 deviating genes are in the 21q22.1 region where Wu et al. observed A/B compartment switches in T21 fetal brain high throughput chromosome conformation capture (Hi-C) [23]. In either direction, genetic effects not captured by common *cis*-eQTLs, such as rare regulatory variants, splicing QTLs (sQTLs), or *trans*-eQTLs, may also contribute. Integrating evidence from additional omic profiles would enable the direct testing of multiple alternative hypotheses.

Our study has several limitations. The eQTLs we tested were mapped in euploid GTEx whole blood. GTEx mapped them in bulk tissue, which dilutes cell-type-specific effects [24]. Donovan et al. found that whole-blood cell-type composition differed in subpopulations of this cohort [4]. Cell-type-specific eQTLs, from single-cell data such as OneK1K [25], might be detected where bulk eQTLs were not. Moreover, ancestry may affect the observed eQTLs. Ancestry structure in the cohort was handled through covariates rather than genotype principal components (Supplement B).

The covariates, chosen for their association with karyotype, also affected which genes deviate. The unadjusted and adjusted analyses share 6 deviating genes. The composition adjustment covers leukocytes only, so a gene expressed in erythroid or platelet lineages, such as *KCNE1*, could deviate through a composition difference the model cannot see. *OLIG2* and *BACE2* deviate in all tested models. Even in the adjusted model, 9 of the 13 deviating genes lie between 33% and 50% from the expected dosage, within about one standard error of the threshold, and a larger cohort would resolve them.

These results are relevant for the genomic hypothesis of embryonic selection. About 80% of T21 conceptions are lost before birth [7], and live-born people with DS have fewer deleterious variants on HSA21 than expected [8]. If survival to term favored embryos whose common *cis*-regulatory alleles dampen dosage-sensitive HSA21 genes, those alleles would occur at shifted frequencies in live-born people with T21. We saw no such shift: across the 2,076 tested *cis* variants, the T21 frequencies did not differ systematically from those of GTEx donors. Most eQTLs mapped in healthy adult tissue have little phenotypic impact [26], though such has not yet been tested in people with DS. Future work should consider rare variants, splicing QTLs, cell-type-specific eQTLs from single-cell data, effects across many loci, interactions between loci, and the representation in people with DS and co-occurring conditions.

In the largest cohort to date with matched transcriptomes and ploidy-aware genotypes, common *cis*-eQTLs act on HSA21 genes in people with T21 as they do in D21 blood. Common eQTLs do not explain deviations in gene expression from the expected fold change since similar allele frequencies were observed in T21 and D21. Therefore, deviations must arise from other mechanisms related to the trisomy itself.

## Data and Code Availability

The demographics and clinical data for research participants in the HTP study are available on both the Synapse data sharing platform (https://doi.org/10.7303/syn31488784) and the INCLUDE Data Hub (https://portal.includedcc.org/). Whole-blood transcriptome data for research participants are available through Synapse (https://doi.org/10.7303/syn31488780), the INCLUDE Data Hub, and Gene Expression Omnibus (GSE190125). Mass cytometry data for 380+ research participants are available in Synapse (https://doi.org/10.7303/syn31488783). Whole genome sequencing data are available under controlled access in the NIH database of Genotypes and Phenotypes (dbGaP) under accession phs002981.v2.p1 and require a Data Access Request approved by the NHLBI Data Access Committee (https://www.ncbi.nlm.nih.gov/projects/gap/cgi-bin/study.cgi?study_id=phs002981.v2.p1). Whole-blood *cis*-eQTL data were obtained from the GTEx Portal (release v10; https://gtexportal.org/).

All original code for ploidy-aware variant calling on HSA21 has been deposited at https://github.com/pivlab/dosage_comp and is publicly available as of the date of publication. Code for the ploidy-aware differential expression and common eQTL comparisons is available at https://github.com/greenelab/T21-eQTL.

## Acknowledgements

This work was funded by NIH grant R01 HD109765.

The results analyzed and published here use data generated by the INCLUDE Project (dbGaP accession phs002981.v2.p1), and were accessed from the INCLUDE Data Hub and/or dbGaP (www.ncbi.nlm.nih.gov/gap). We thank the Human Trisome Project investigators at the Linda Crnic Institute for Down Syndrome, who submitted these data. Moreover, we thank the study participants and their families. The GTEx Project was supported by the Common Fund of the Office of the Director of the National Institutes of Health, and by the National Cancer Institute, the National Human Genome Research Institute, the National Heart, Lung, and Blood Institute, the National Institute on Drug Abuse, the National Institute of Mental Health, and the National Institute of Neurological Disorders and Stroke. The data used for the analyses described in this manuscript were obtained from the GTEx Portal (release v10) on May 4, 2026.

## Declaration of generative AI and AI-assisted technologies

This project used the AI assistant Claude, developed by Anthropic, during the development of the analysis code. Its assistance included generating initial code for the analyses and figures, and improving the code documentation. Additionally, AI tools were used to critique drafts of this article and to identify opportunities to better communicate with a broader audience. The authors reviewed, tested, and validated all AI-generated content and take full responsibility for the content of this article.

