## Supplement for "Common *cis*-eQTLs do not explain chromosome 21 gene expression deviations from the ploidy expectation in Trisomy 21"


- **Lucas A. Gillenwater** ✉
  0000-0003-3158-9682
  Department of Biomedical Informatics, University of Colorado Anschutz, Aurora, CO

- **Mary A. Allen**
  0000-0001-7490-0165
  BioFrontiers Institute, University of Colorado Boulder, Boulder, CO

- **Milton Pividori**
  0000-0002-3035-4403
  Department of Biomedical Informatics, University of Colorado Anschutz, Aurora, CO

- **Casey S. Greene**
  0000-0001-8713-9213
  Department of Biomedical Informatics, University of Colorado Anschutz, Aurora, CO

- **Marc Subirana-Granés** ✉
  0000-0003-3934-839X
  Department of Biomedical Informatics, University of Colorado Anschutz, Aurora, CO

### Supplement

#### Supplement A: Evidence for covariate adjustment

Before attributing a gene's deviation from the 1.5-fold expectation to HSA21 dosage rather than age, BMI, or blood-cell composition, we confirmed, in the full cohort, that: (1) several clinical and demographic factors differ significantly by karyotype (age p = 0.033, BMI p < 0.0001; sex did not differ as strongly (p=0.0982) though still differed); (2) directly measured blood-cell-type composition is associated with expression of thousands of genes genome-wide; and (3) the 9 mosaic-DS participants, whose cells do not all carry the extra chromosome, show a smaller chromosome-21 dosage effect (mean composite HSA21 index ≈ 1.07x) than participants with full trisomy (≈1.35–1.45x), a sanity check that the dosage measurement itself behaves as expected. These findings motivate adjusting for age, sex, BMI, sample source, and cell-type composition in the primary analysis (Figure S1).

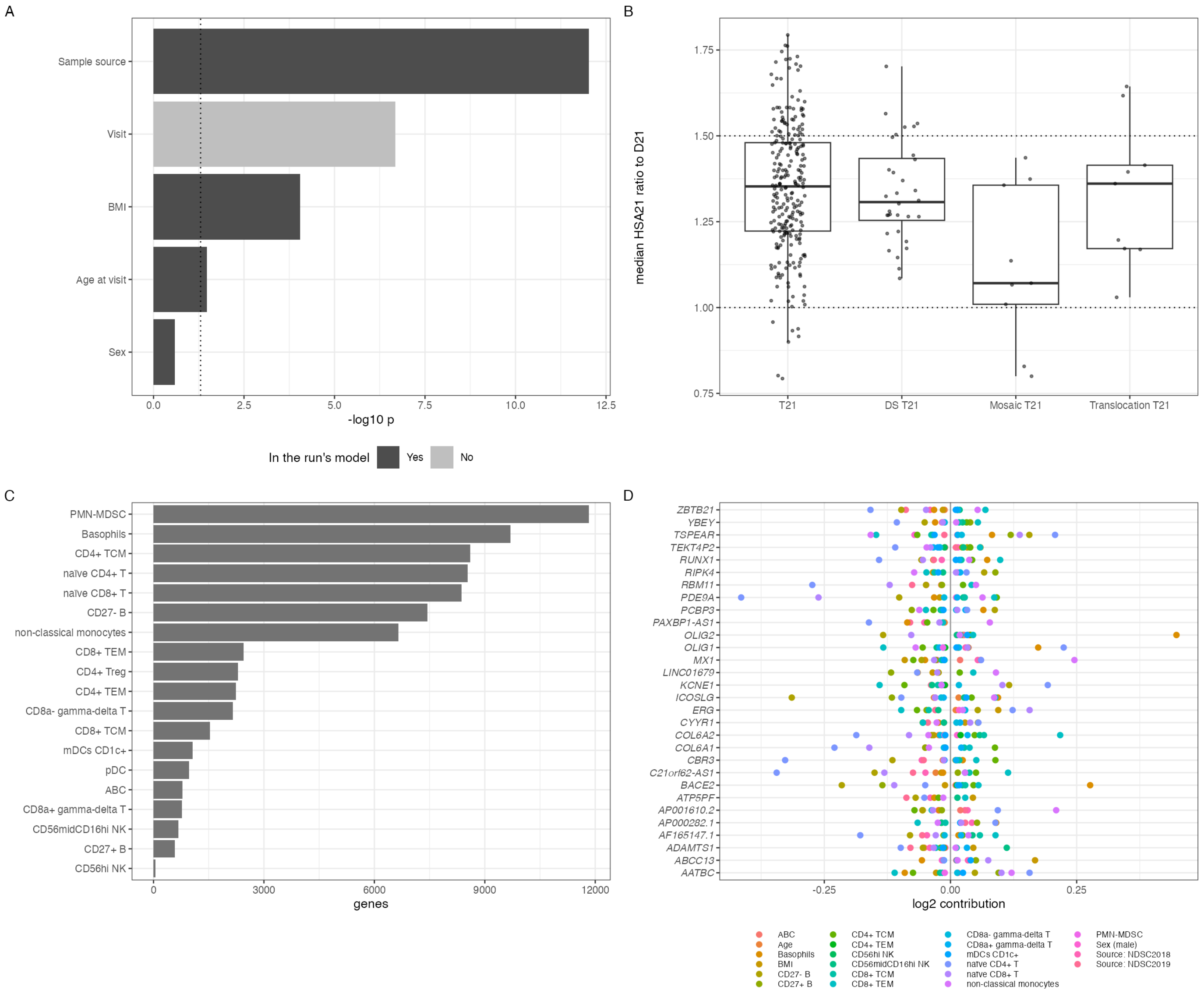


**Figure S1:** Covariate evidence for the adjusted analysis. (A) Association of each candidate covariate with karyotype ($-\log_{10}$ p), colored by whether it entered the adjusted model; the dotted line marks p = 0.05. (B) HSA21 expression index (median ratio of HSA21 gene expression to D21) by karyotype subtype as recorded in the INCLUDE Data Hub (DS T21, Down syndrome with the subtype unspecified), with reference lines at 1 and 1.5; mosaic participants sit near 1. (C) Number of genes associated with each CyTOF cell fraction within T21 at 5% FDR. (D) For each gene that deviates in either analysis, the contribution of each covariate to the shift in $\log_2$ fold change between the unadjusted and adjusted analyses, colored by covariate.

### Supplement B: Genotype structure in the within-T21 eQTL tests

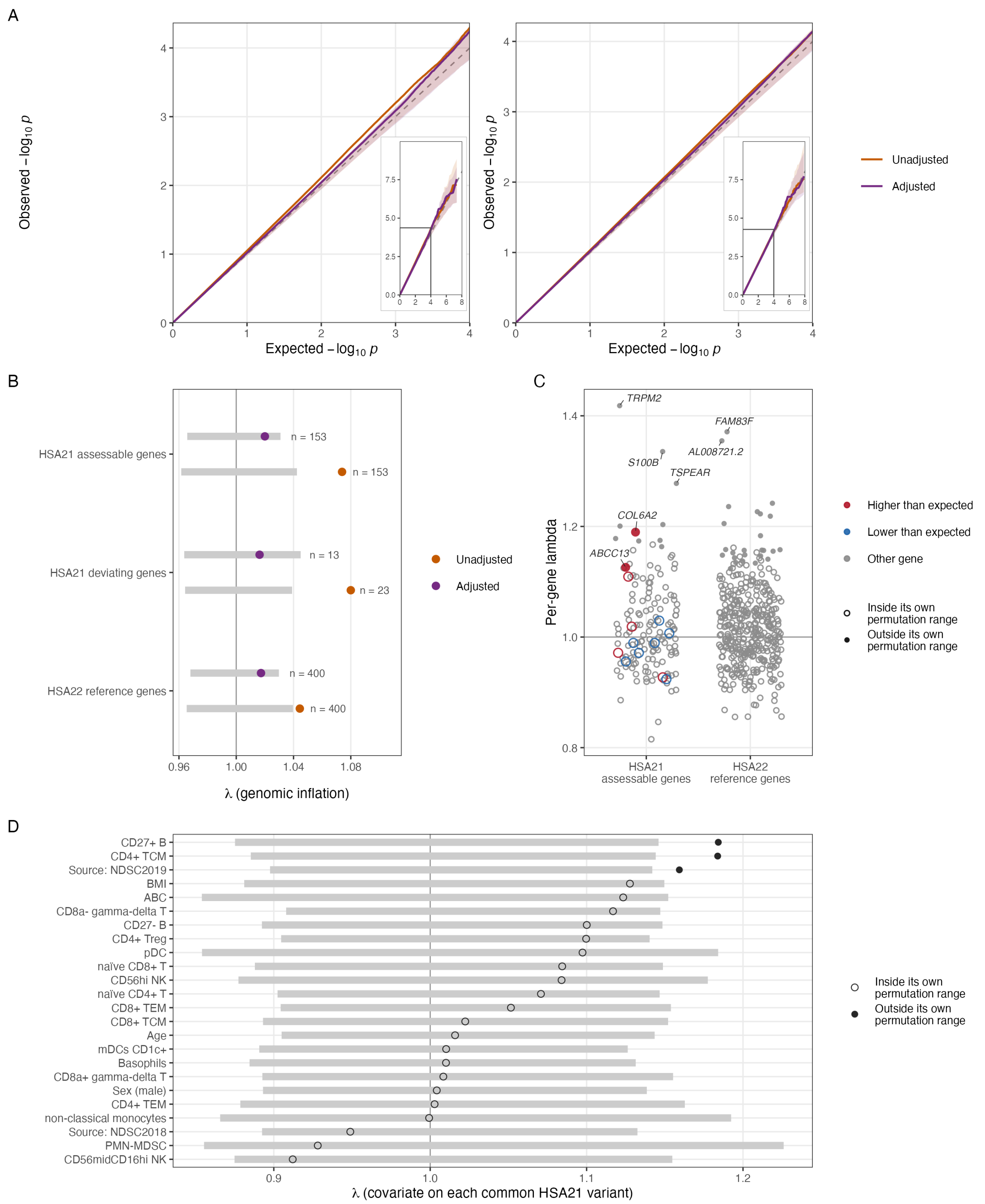


**Figure S2:** Genotype structure in the within-T21 tests. (A) Quantile–quantile plots of T21 expression regressed on common HSA21 long-arm variants at least 5 Mb from each gene, for the assessable HSA21 genes (left) and for HSA22 reference genes (right), without covariate adjustment (unadjusted) and with the covariates regressed out of expression and genotype (adjusted); shaded, 95% range over 200 participant permutations. Each plot shows expected $-\log_{10} p$ up to 4, where most tests lie; the inset shows the full range with that window outlined. (B) Genomic inflation factor λ per gene set with its permutation 95% range. (C) Per-gene λ in the adjusted analysis, HSA21 assessable and HSA22 reference genes, with deviating genes colored by direction (red, higher than expected; blue, lower than expected); filled points lie outside the gene's own permutation 95% range. The five genes with the highest λ and the deviating genes outside their own range are labeled. (D) Each covariate of the adjusted analysis regressed on every common HSA21 long-arm variant; filled points lie outside their own permutation 95% range.

The *cis*-eQTL model does not include genotype principal components, so we measured genotype structure in the within-T21 tests directly. For each of the 153 assessable HSA21 genes (not repeat-flagged, base mean ≥ 30), T21 expression was regressed on every common variant on the HSA21 long arm (minor allele frequency ≥ 0.05 and call rate ≥ 0.95 among the T21 participants analyzed; 97,438 variants) at least 5 Mb from the gene's transcription start site, one ordinary least-squares regression per gene–variant pair (11.1 million pairs). At that distance a variant cannot tag the gene's own *cis* locus, so in the absence of structure the p-values are uniform and the genomic inflation factor λ is 1. Variants on the short arm and around the centromere were set aside: GRCh38 places false duplications of 21q22.3 there [1,2], so a variant's coordinate can put a long-arm gene's own *cis* signal more than 5 Mb away (for *GATD3A*, *PWP2*, *SIK1*, and *ICOSLG*, down to $p \approx 10^{-46}$); including them changes λ by less than 0.02. Two references were used to read λ. The same regressions were run for 400 expressed HSA22 genes: population structure would inflate their tests as much as those of HSA21 genes, whereas a signal tied to HSA21 copy number would not. Participant labels were also permuted 200 times, jointly across genes, which keeps the correlation between genes and the linkage disequilibrium between variants; the spread of λ over these permutations is its sampling range for these correlated tests, about ±0.04 for a gene set.

Without covariate adjustment (302 T21 participants), λ was 1.074 for HSA21 genes and 1.044 for HSA22 genes, both above their permutation ranges, and 26 HSA21 genes lay outside their own permutation range, against a permutation median of 3 (Figure S2A, B). In the primary analysis (274 T21 participants), with age, sex, BMI, sample source, and cell-type composition regressed out of both expression and genotype dosage as in the *cis*-eQTL test, λ was 1.020 for HSA21 genes (permutation range 0.966–1.031), 1.016 for the 13 deviating genes, and 1.017 for HSA22 genes, all within their permutation ranges. Three of the 24 adjustment covariates were themselves associated with HSA21 genotype across the chromosome, against a permutation median of 0 (Figure S2D): recruitment at NDSC2019 and the CD27+ B-cell and CD4+ central memory T-cell fractions. The adjusted tests are therefore not inflated beyond their permutation range at the level of the gene set, but the adjustment reaches that through covariates that track genotype structure rather than through a direct measure of ancestry, and some gene-level structure may remain: 11 HSA21 genes lay outside their own permutation range against a permutation median of 3 ($p = 0.04$), among them two deviating genes, *COL6A2* and *ABCC13* (permutation $p = 0.020$ and 0.025; Figure S2C).

#### Supplement C: Comparison across covariate models

Running the identical pipeline without covariate adjustment and without excluding mosaic participants (397 participants: 302 T21, 95 D21) identified 23 deviating genes (10 higher, 13 lower) rather than 13, and detected a *cis*-eQTL for 14 of 20 testable genes rather than 8 of 10. Of the 13 genes that deviate in the primary (adjusted) analysis, 6 also deviate in the unadjusted analysis (*OLIG2*, *PCBP3*, *BACE2*, *YBEY*, *COL6A2*, *ABCC13*); the other 7 deviate only once age, BMI, and cell-type composition are accounted for (*KCNE1*, *OLIG1*, *RUNX1*, *AP000282.1*, *CBR3*, *ADAMTS1*, *ATP5PF*). Of the 23 genes that deviate in the unadjusted analysis, 17 drop out once adjusted — though for 11 of those, the effect size is similar in the adjusted model and the gene simply falls short of the stricter significance threshold in the smaller, more heavily parameterized model, rather than the underlying effect disappearing. This comparison is the basis for reporting the adjusted analysis as primary while retaining the unadjusted analysis for transparency (Figure S3).

The adjusted model assumes that each cell fraction affects expression in the same way in T21 and D21. An interaction model, with a karyotype × cell-fraction interaction, raised the standard error of the T21 effect by a median of 1.76-fold for every HSA21 gene. We compared effect estimates rather than significance across the three models (Table S3). Four genes deviated under both the adjusted and the interaction models, all lower than expected: *OLIG2*, *BACE2*, *OLIG1*, and *RUNX1*. *OLIG2* and *BACE2* also deviated without adjustment. *OLIG1* and *RUNX1* were lower than expected without adjustment as well, but below the magnitude cut. Five genes kept similar estimates under the interaction model but lost significance: *YBEY*, *ADAMTS1*, *ABCC13*, *AP000282.1*, and *COL6A2*, which also fell just below the magnitude cut. The estimates for *KCNE1* and *PCBP3* shrank by about a third. The estimates for *CBR3* and *ATP5PF* disappeared, and neither gene deviated without adjustment, so their deviations depend on the assumption that composition relates to expression alike in both groups. None of the six genes higher than expected deviated under the interaction model, and no gene outside the 13 began to deviate. *OLIG1*, *OLIG2*, and *AP000282.1* lie within 60 kb of each other and their expression is correlated (Supplement E), so the four most robust genes may reflect three independent deviations.

**Table S3:** Ploidy-corrected $\log_2$ fold change (95% CI) of the 13 deviating genes under the unadjusted, adjusted (primary), and karyotype × cell-fraction interaction models. Bold estimates deviate in that model (adjusted $p < 0.01$ and a fold change of at least 1.33 in either direction). Genes are ordered from the most to the least robust.

| Gene | Direction | Unadjusted | Adjusted | Interaction |
|---|---|---|---|---|
| *OLIG2* | Lower | **−1.19 (−1.44, −0.94)** | **−2.18 (−2.63, −1.72)** | **−2.65 (−3.43, −1.88)** |
| *BACE2* | Lower | **−0.43 (−0.56, −0.30)** | **−0.58 (−0.79, −0.36)** | **−0.89 (−1.27, −0.51)** |
| *OLIG1* | Lower | −0.36 (−0.50, −0.23) | **−0.76 (−0.99, −0.52)** | **−0.97 (−1.37, −0.56)** |
| *RUNX1* | Lower | −0.31 (−0.37, −0.26) | **−0.43 (−0.53, −0.33)** | **−0.42 (−0.60, −0.25)** |
| *YBEY* | Higher | **0.50 (0.40, 0.59)** | **0.47 (0.30, 0.64)** | 0.49 (0.18, 0.79) |
| *ADAMTS1* | Higher | 0.21 (0.02, 0.39) | **0.48 (0.22, 0.73)** | 0.51 (0.06, 0.97) |
| *ABCC13* | Higher | **1.45 (1.17, 1.73)** | **1.01 (0.49, 1.52)** | 0.83 (−0.08, 1.73) |
| *AP000282.1* | Lower | −0.27 (−0.43, −0.11) | **−0.70 (−0.99, −0.42)** | −0.87 (−1.37, −0.37) |
| *COL6A2* | Higher | **0.56 (0.41, 0.71)** | **0.42 (0.19, 0.64)** | 0.38 (−0.01, 0.77) |
| *KCNE1* | Lower | −0.38 (−0.52, −0.24) | **−0.50 (−0.72, −0.28)** | −0.36 (−0.74, 0.03) |
| *PCBP3* | Lower | **−0.43 (−0.57, −0.29)** | **−0.50 (−0.76, −0.25)** | −0.31 (−0.75, 0.14) |
| *CBR3* | Higher | 0.10 (−0.05, 0.25) | **0.53 (0.27, 0.78)** | 0.00 (−0.43, 0.44) |
| *ATP5PF* | Higher | 0.27 (0.13, 0.41) | **0.48 (0.22, 0.74)** | 0.19 (−0.24, 0.62) |

**Table S4:** Classification of all 318 targeted HSA21 genes in the primary (adjusted) analysis.

| Classification | Subgroup | Genes |
|---|---|---:|
| Expected dosage | | 129 |
| Outside the expectation (24) | Did not reach significance | 11 |
| | Higher than expected | 6 |
| | Lower than expected | 7 |
| Not assessable (165) | Low expression | 153 |
| | High repeats | 12 |
| Total | | 318 |

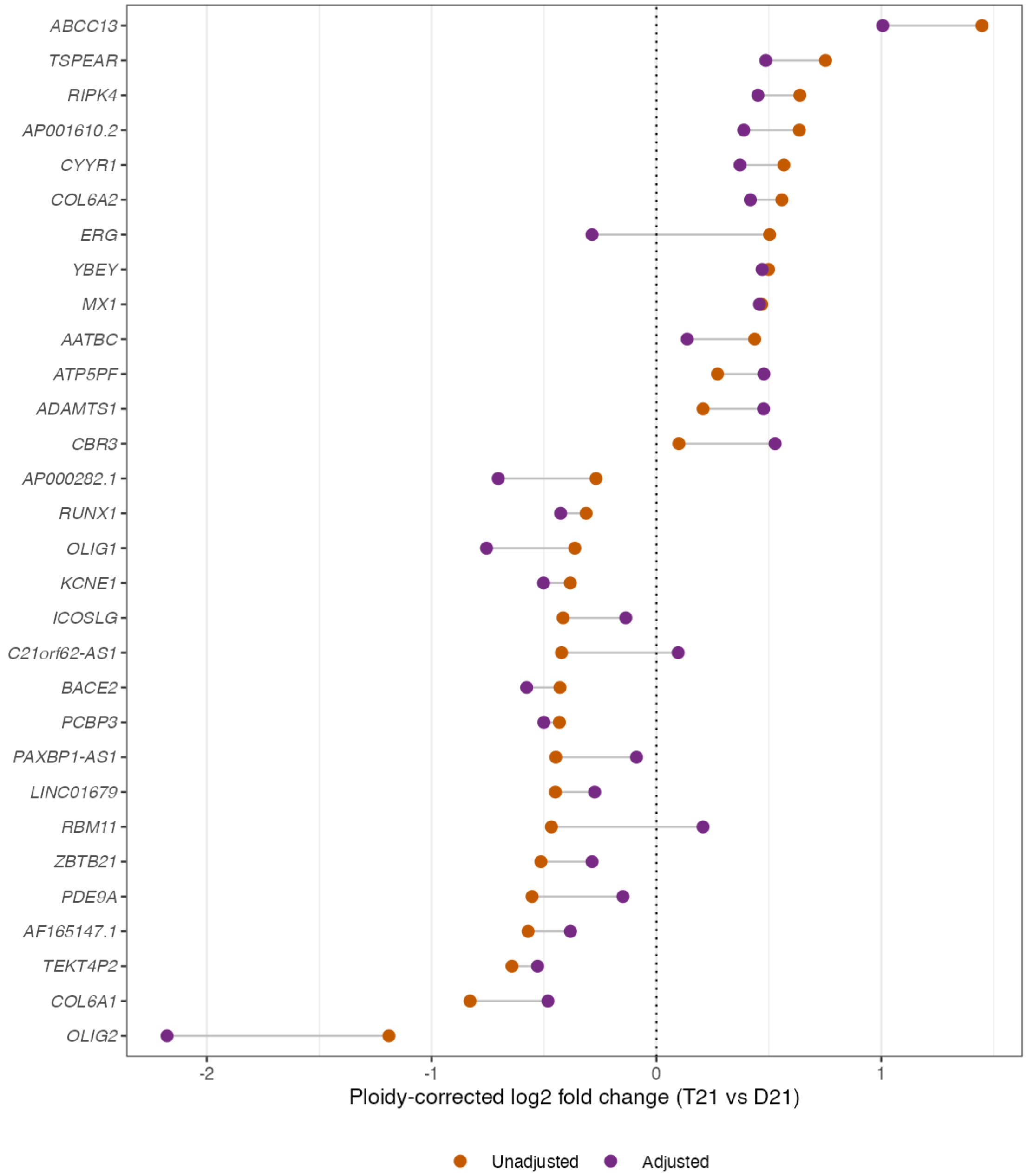


**Figure S3:** Comparison of the adjusted (primary) and unadjusted analyses. One row per gene that deviates in either analysis, showing its ploidy-corrected $\log_2$ fold change under the unadjusted (orange) and adjusted (purple) analyses, joined by a line; the dotted line marks zero.

### Supplement D: Whole genome differential expression

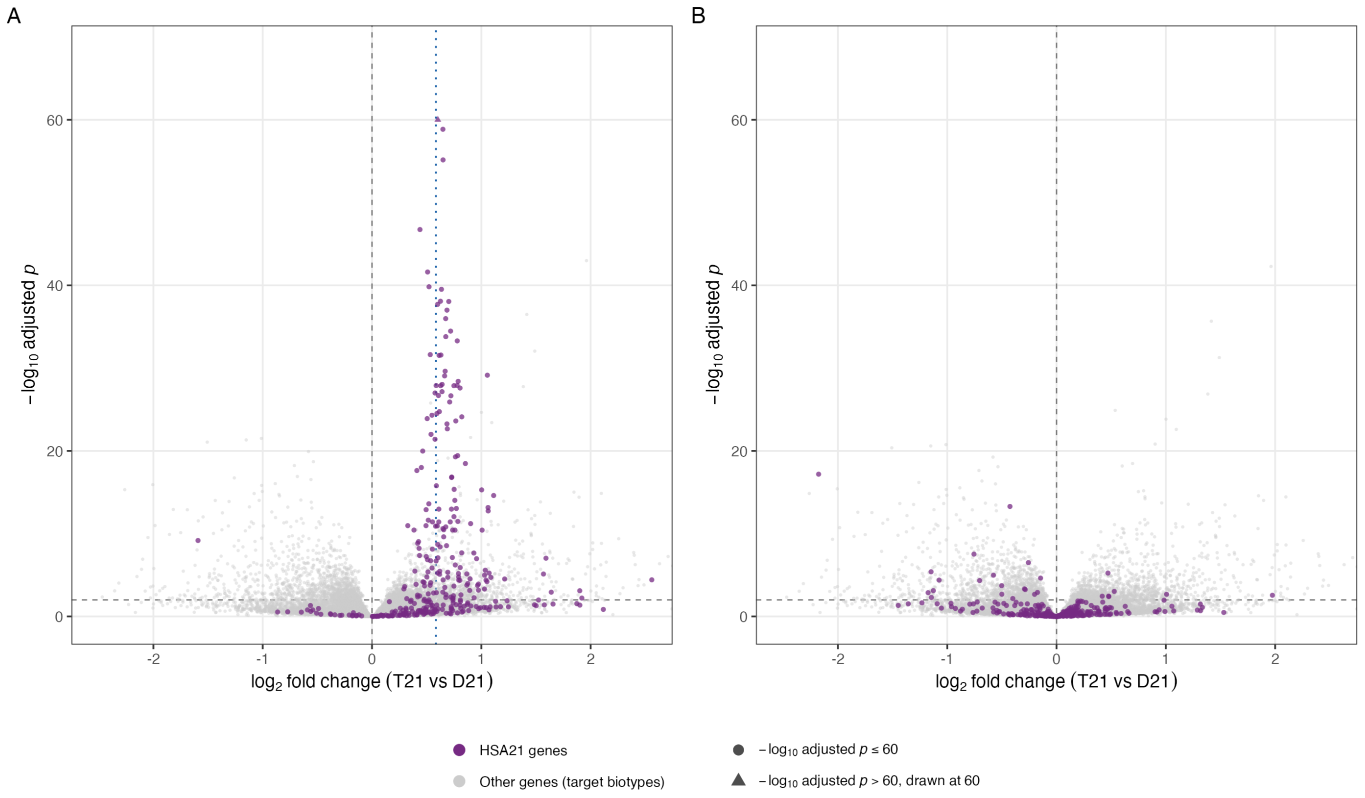


**Figure S4:** Differential expression of all genes of the targeted biotypes (protein-coding, lncRNA, pseudogene) genome-wide, before (A) and after (B) ploidy correction, with HSA21 genes highlighted and all other genes in grey. The dotted line in A marks the 1.5-fold expectation; before correction the HSA21 genes cluster around it and are almost uniformly significant, and after correction they center on zero.

### Supplement E: Expression of the HSA21 genes in blood

We compared HSA21 gene expression in GTEx v10 whole blood and other tissues. The deviating genes' whole-blood expression was lower than that of the expected-dosage genes, but not significantly (median 1.3 vs 3.1 TPM, Wilcoxon p = 0.34; Figure S5A).

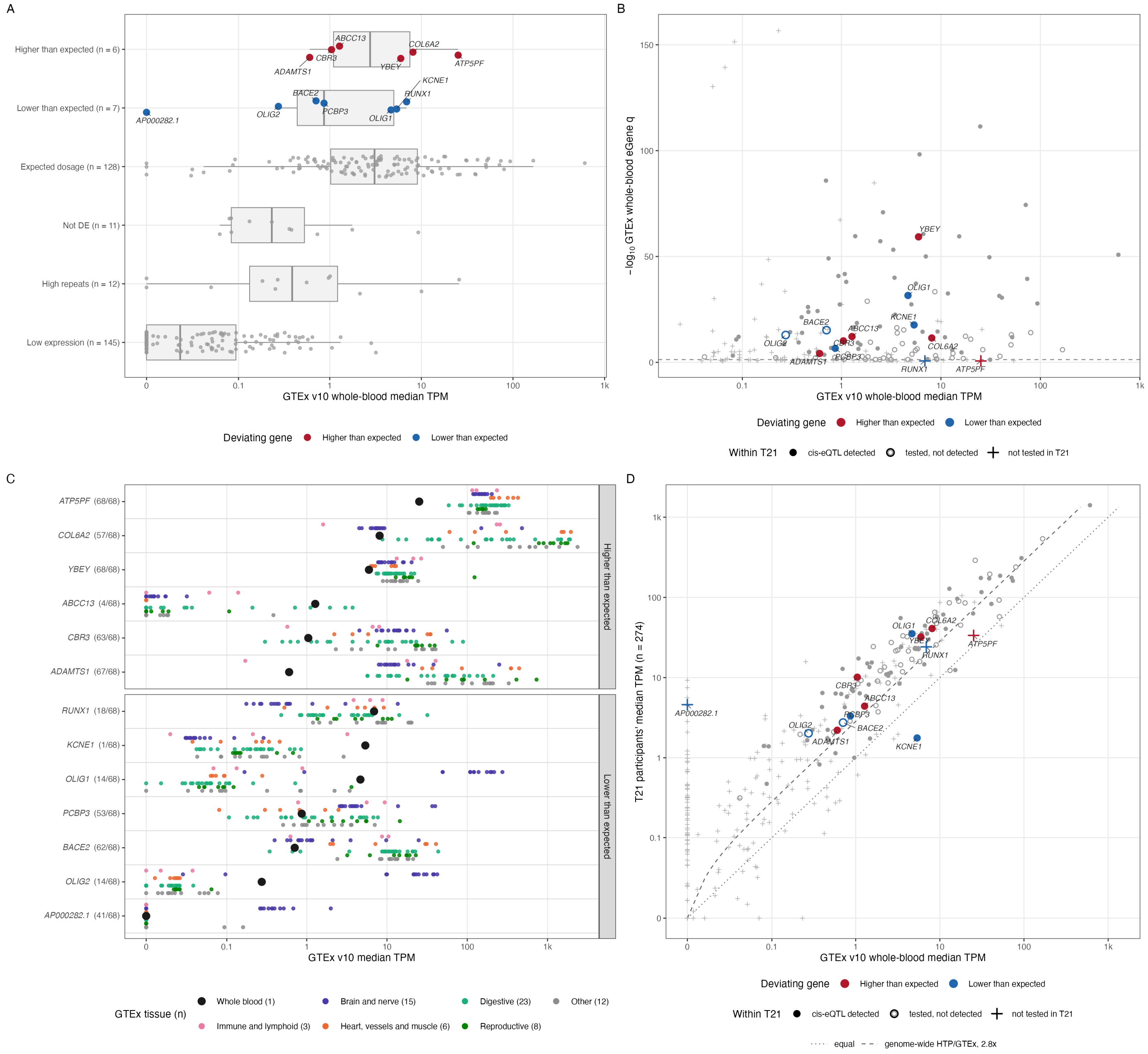


**Figure S5:** Expression of the HSA21 genes in blood. (A) GTEx v10 whole-blood median TPM of the classified HSA21 genes by classification. Deviating genes are colored by direction. (B) GTEx whole-blood eGene q-value ($-\log_{10}$) against whole-blood TPM for the 211 HSA21 genes that GTEx tested in whole blood. Filled points have a *cis*-eQTL detected in T21, open points were tested in T21 without detection, and crosses were not tested in T21 (102 genes); deviating genes are colored by direction and the dashed line marks q = 0.05. (C) Median TPM of each deviating gene in the 68 GTEx tissues. Whole blood (large point) is on the gene's line, and the other tissues are grouped by organ system below it. Labels give the rank of whole blood among the 68 tissues. (D) Median TPM in the 274 T21 participants against GTEx whole-blood TPM for the 309 classified HSA21 genes in GTEx, with point shapes as in B. The dotted line marks equality, and the dashed line marks the genome-wide HTP/GTEx offset (2.8-fold).

Expression did not predict eQTL detection (Figure S5B). Among the 211 HSA21 genes GTEx tested in whole blood, eGenes and non-eGenes had similar expression (median 1.28 vs 1.18 TPM, p = 0.36). Genes with a *cis*-eQTL detected in T21 were, if anything, less expressed than those without (median 2.4 vs 3.9 TPM, p = 0.13).

Whole blood ranks low among the GTEx tissues for most genes (median rank 64 of 68 for the expected-dosage genes; Figure S5C). Expression in the T21 participants tracked GTEx closely (Spearman 0.87 across the 309 HSA21 genes; Figure S5D). The HTP values were 2.8-fold higher overall. Part of this offset may come from globin depletion: the three main hemoglobin genes take 0.4% of HTP TPM, against 30% in GTEx whole blood.

*AP000282.1* has zero expression in GTEx whole blood but a median of 4.6 TPM in the T21 participants. Its first exon lies inside the single exon of *OLIG1*, on the opposite strand. GTEx libraries are unstranded, so GTEx removes exon regions shared between genes before counting [3]. The HTP libraries are stranded and were counted by strand, so the T21 counts belong to *AP000282.1* itself. *AP000282.1* is correlated with *OLIG1* (Spearman 0.67 in T21 after covariate adjustment). *OLIG1*, *OLIG2*, and *AP000282.1* may deviate as one locus rather than independently.
